# The Neuron Specific Formin Delphilin Nucleates Actin Filaments but Does Not Enhance Elongation

**DOI:** 10.1101/151191

**Authors:** William T. Silkworth, Kristina L. Kunes, Grace C. Nickel, Martin L. Phillips, Margot E. Quinlan, Christina L. Vizcarra

**Author notes:** co-corresponding authors: clv and meq.

## Abstract

The formin Delphilin binds the glutamate receptor, GluRδ2, in dendritic spines of Purkinje cells. Both proteins play a role in learning. To understand how Delphilin functions in neurons, we studied the actin assembly properties of this formin. Formins have a conserved formin homology 2 domain, which nucleates and remains associated with the fast growing end of actin filaments, influencing filament growth with input from the adjacent formin homology 1 domain. The strength of nucleation and elongation varies widely across formins. Additionally, most formins have conserved domains that regulate actin assembly through an intramolecular interaction. Delphilin is distinct from other formins in several ways: its expression is limited to Purkinje cells; it lacks autoinhibitory domains; its formin homology 1 domains has minimal proline-rich sequence. We found that Delphilin is an actin nucleator that does not accelerate elongation, although it binds tightly to the barbed end of filaments. In addition, Delphilin exhibits a preference for actin isoforms, which has not been described or systematically studied in other formins. Finally, Delphilin is the first formin studied that is not regulated by intramolecular interactions. We speculate how the activity we observe is consistent with its localization in the small dendritic spines.

Abbreviations: FH1, formin homology 1; FH2, formin homology 2; DID, diaphanous inhibitory domain; DAD, diaphanous autoinhibitory domain; LTD, long term depression; PSD, post synaptic density; PDZ, <u>p</u>ost synaptic density protein (PSD95), <u>D</u>rosophila disc large tumor suppressor (Dlg1), and <u>z</u>onula occludens-1 protein (zo-1).

## Introduction

Dendritic spines are post-synaptic structures essential for learning (Nimchinsky *et al.*, 2002). Defects in spine morphology and/or number are linked to a number of disorders, including Autism Spectrum Disorder, Schizophrenia, and Alzheimers Disease (Penzes *et al.*, 2011). It is noteworthy that symptoms of the three disorders listed here present at very different ages. Thus development and maintenance of dendritic spines are both important subjects. Spine morphology is dynamic and reflects synaptic activity levels. Spines are commonly described as mushroom shaped structures, with a head and neck, though the morphology varies from filopodia-like to mushroom shaped (Peters and Kaiserman-Abramof, 1970; Harris *et al.*, 1992). The head contains the so-called postsynaptic density (PSD), a collection of 100s of proteins including receptors, signaling proteins, and cytoskeletal elements (Walikonis *et al.*, 2000). The actin cytoskeleton is essential for formation, maintenance, and dynamic rearrangement of dendritic spines (Hlushchenko *et al.*, 2016). It follows that many actin binding proteins are found in spines.

Actin nucleators help build new filaments and structures in a manner related to their respective nucleation mechanisms. The Arp2/3 complex, several formins, and, perhaps, tandem monomer binding nucleators all contribute to dendritic spine morphology and dynamics. The nucleators build distinct portions of the spines and respond to different events (Hotulainen *et al.*, 2009; Hlushchenko *et al.*, 2016). The formin, Delphilin, is localized exclusively in neurons and primarily in cerebellar dendritic spines; however, it is not required for spine morphology and its role remains unclear (Takeuchi *et al.*, 2008). Delphilin was first identified as a binding partner for the glutamate receptor, GluRδ2 (Miyagi *et al.*, 2002). The GluRδ2 receptor is critical to synaptic plasticity and motor learning. The GluRδ2 receptor is also a bit of a mystery, with no known ligand or apparent electrical activity. Delphilin and GluRδ2 interact in Purkinje cells at synapses with parallel fibers (Yamashita *et al.*, 2005). Loss of GluRδ2 results in disrupted synapse formation and loss of long term depression (LTD), a case when synaptic connections lose efficiency (Kashiwabuchi *et al.*, 1995). While Delphilin is not required for spine formation, its loss does result in facilitation of LTD, consistent with its known interaction with GluRδ2 (Takeuchi *et al.*, 2008).

Mammals have 15 formins which represent seven of nine formin families (Higgs and Peterson, 2005; Pruyne, 2016). It is distinct from most formins in that it does not contain a Diaphanous Inhibitory or Autoinhibitory Domains (DID and DAD). Formins in six of the nine formin families are regulated by intramolecular interactions between DID and DAD domains. Fmn-family formins do not have DID and DAD domains but are autoinhibited by an analogous intramolecular interaction (Bor *et al.*, 2012). Instead of a DID domain, the three Delphilin splice variants have at least one PSD-95/Discs large/ZO-1 (PDZ) domain at the N-terminus (one has two) (Yamashita *et al.*, 2005). PDZ domains are common to PSD proteins and, logically, the Delphilin-PDZ domain binds directly to the C-terminal tail of GluRδ2 (Miyagi *et al.*, 2002). The Delphilin tail, where the DAD domain would be, is essentially absent.

It is easy to speculate that Delphilin’s actin assembly activities are essential to its function. However, Delphilin-family formins have not been extensively characterized. Therefore, to determine if actin assembly in response to synaptic activity could be part of Delphilin’s role, we compared it to other well studied formins, in vitro. Formins are identified based on conserved formin homology 1 and 2 (FH1 and FH2) domains, which mediate actin nucleation and processive elongation, in most cases. We found that a construct containing the FH1 and FH2 domains of Delphilin assembles actin in a manner similar to the previously characterized formin, Cdc12 (Kovar *et al.*, 2003). In the absence of profilin, new filaments only grow from their pointed ends and in the presence of profilin the barbed ends grow. However, the elongation rate is not accelerated above actin alone. Additionally, we found that, unlike other studied formins, Delphilin does not appear to be autoinhibited.

## Results

### Delphilin is an actin assembly factor

To understand the effect Delphilin has on actin assembly we purified the C-terminal halves of both mouse and human Delphilin (mDelFFC and hDelFFC), including both the FH1 and FH2 domains through to their C-termini (Figures 1A and S1A). We tested both formins in pyrene-actin assembly assays. Addition of either DelFFC construct results in a dose-dependent increase in actin assembly (Figures 1B and S2A). In this context, mDelFFC is ~5 fold more potent than hDelFFC. The minimum domain required for actin assembly by most formins is the FH2 domain (Pruyne *et al.*, 2002; Li and Higgs, 2003; Goode and Eck, 2007). To verify that the Delphilin FH1 domain is dispensable for actin assembly in the absence of profilin we tested mDelFC (Figure 1A). We found that mDelFC assembles actin filaments, albeit ~15-fold slower than mDelFFC (Figure 1C). The majority of free actin monomers within cells are bound by the actin-binding protein, profilin. Therefore, to test mDelFFC under more physiologically relevant conditions, we added profilin to the pyrene-actin assembly assays. Under these conditions, actin assembly was markedly decreased but we still observed dose-dependent stimulation of actin polymerization (Figure S2B). These results confirm that Delphilins are capable of enhancing actin assembly, as predicted based on primary sequence.

**Figure 1.**
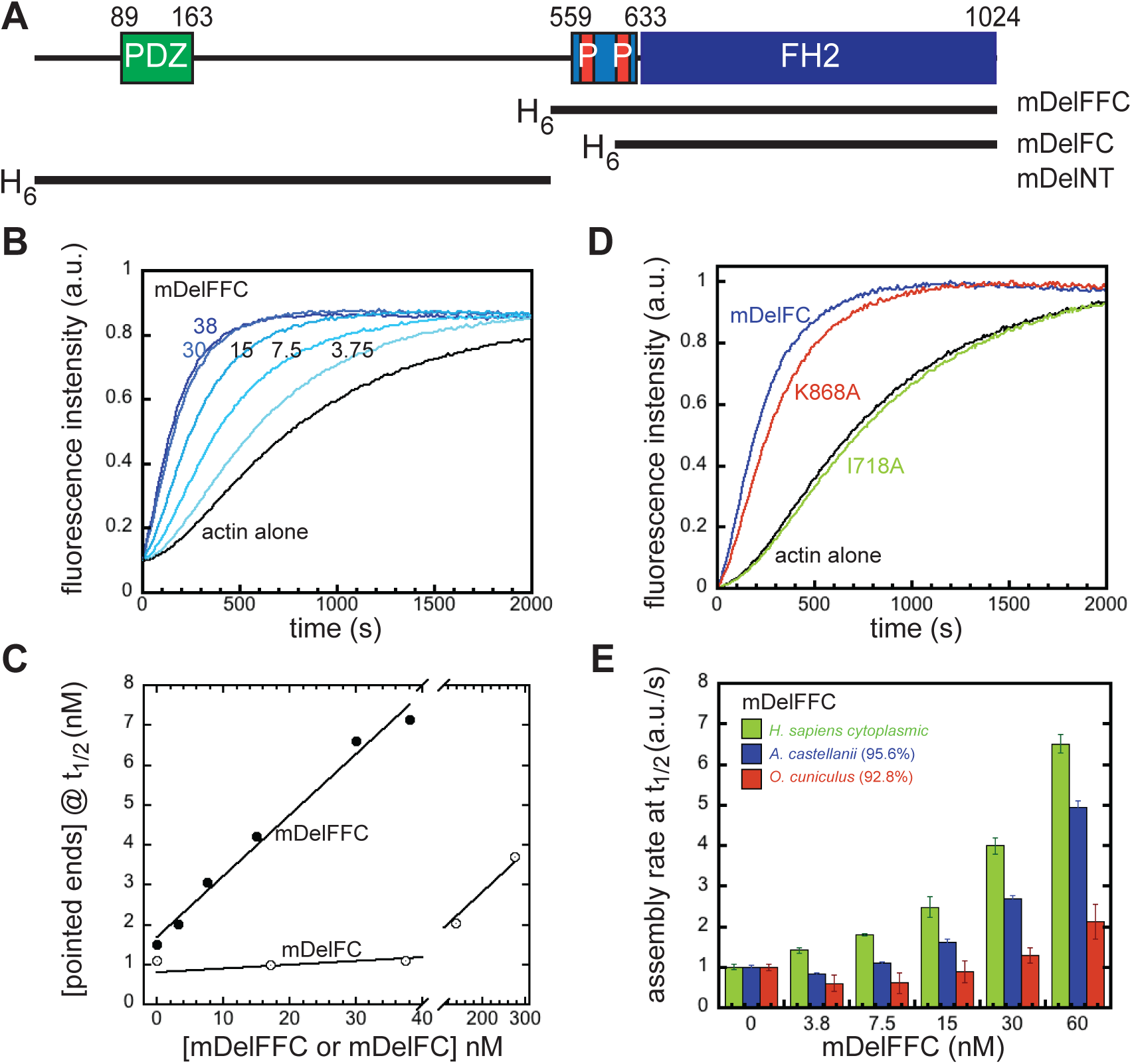
Delphilin is an actin assembly factor. A) Domain structure of Delphilin. Green, PDZ domain; light blue, FH1 domain, with red proline rich repeats; blue, FH2 domain. The numbering is based on mouse α-Delphilin (NP_579933.1). Constructs used in this paper are indicated below the diagram. All constructs were N-terminally His-tagged. B) Pyrene actin assembly assay with mDelFFC concentrations as indicated (nM). C) Comparison of actin assembly by mDelFFC and mDelFC. The concentration of filaments was calculated assuming only pointed-end elongation as explained below. D) Comparison of 150 nM wild type mDelFC with point mutations in the same construct (I718A and K868A). E) Comparison of actin assembly rates in pyrene assays with the indicated source of actin. Sequence similarities compared to human cytoplasmic actin are shown in parentheses. Conditions: Acanthamoeba actin (4 μM, 10% pyrene labeled) was used in (B-D). In (E), the source of actin is indicated.

Two highly conserved residues within the FH2 domain, an isoleucine and a lysine, are essential for actin assembly activities in most formins (Xu *et al.*, 2004; Ramabhadran *et al.*, 2012). While mutations in either residue abolish actin assembly in Bni1 (Xu *et al.*, 2004), the analogous mutations have varying effects in other formins (Ramabhadran *et al.*, 2012). We built a homology model of mDelFC based on the crystal structure of hDaam1 (PDB ID 2J1D) and identified the conserved isoleucine and lysine residues, Ile718 and Lys868, respectively. We substituted alanine at these positions in mDelFC and tested the consequences in the pyrene-actin assembly assay. Actin assembly was abolished for all tested concentrations of mDelFC I718A, while mDelFC K868A resulted in only a mild reduction of actin assembly activity (Figure 1D). Taken together these data suggest that Delphilin functions much like other characterized formins.

### Delphilin activity is sensitive to the actin isoform

In the course of our work we noticed a difference in activity, dependent on the actin isoform present. Actin is highly conserved, often sharing >90% sequence identity across species, making this a surprising observation, though not unprecedented among actin binding proteins (Rubenstein, 1990; McCullough *et al.*, 2011). We tested the commonly used actin isoforms from different sources: *A. castellanii* (amoeba actin) and *O. cuniculus* (rabbit skeletal muscle, α-actin), as well as *H. sapiens* (cytoplasmic, 85% β-actin), reasoning that this is the actin that Delphilin is most likely to encounter *in vivo*. Interestingly, both mouse and human DelFFC constructs accelerated actin assembly of human cytoplasmic actin most effectively (Figures 1E and S2A). Amoeba actin, which shares 95.6% sequence identity to cytoplasmic actin was assembled at ~75% the level observed for cytoplasmic actin (Figure 1E). In contrast, Delphilin weakly assembled rabbit skeletal actin (~20%), despite its 92.8% similarity to cytoplasmic actin (Figure 1E). These data indicate that Delphilin is sensitive to actin isoform and nucleates cytoplasmic actin, the actin it would normally encounter, most effectively.

### Delphilin is a nucleator

The kinetics of the pyrene-actin assembly assay reflect both nucleation and elongation of actin filaments. It is particularly important to distinguish these two activities when studying formins, which can modify each to differing degrees. Further, how Delphilin influences these two distinct processes has implications for how this formin functions *in vivo*. To elucidate between nucleation and/or elongation of actin filaments we took advantage of total internal reflection fluorescence (TIRF) microscopy. First, we incubated 2 μM actin alone or in the presence of 1-50 nM mDelFFC for 5 minutes and then stabilized actin filaments with Alexa488-phalloidin. Phalloidin-stabilized filaments were adsorbed to poly-L-lysine coated coverslips and imaged. Confirming that Delphilin is a nucleator, we observed that as concentrations of mDelFFC increase so too do the number of actin filaments (Figure 2A). Compared to actin alone, addition of 50 nM mDelFFC results in greater than 10-fold increase in the number of actin filaments (Figure 2A). We also noted that filaments nucleated by DelFFC were shorter than actin alone filaments (Figure 2A), reminiscent of the yeast formin Cdc12, which caps the barbed end of filaments.

**Figure 2.**
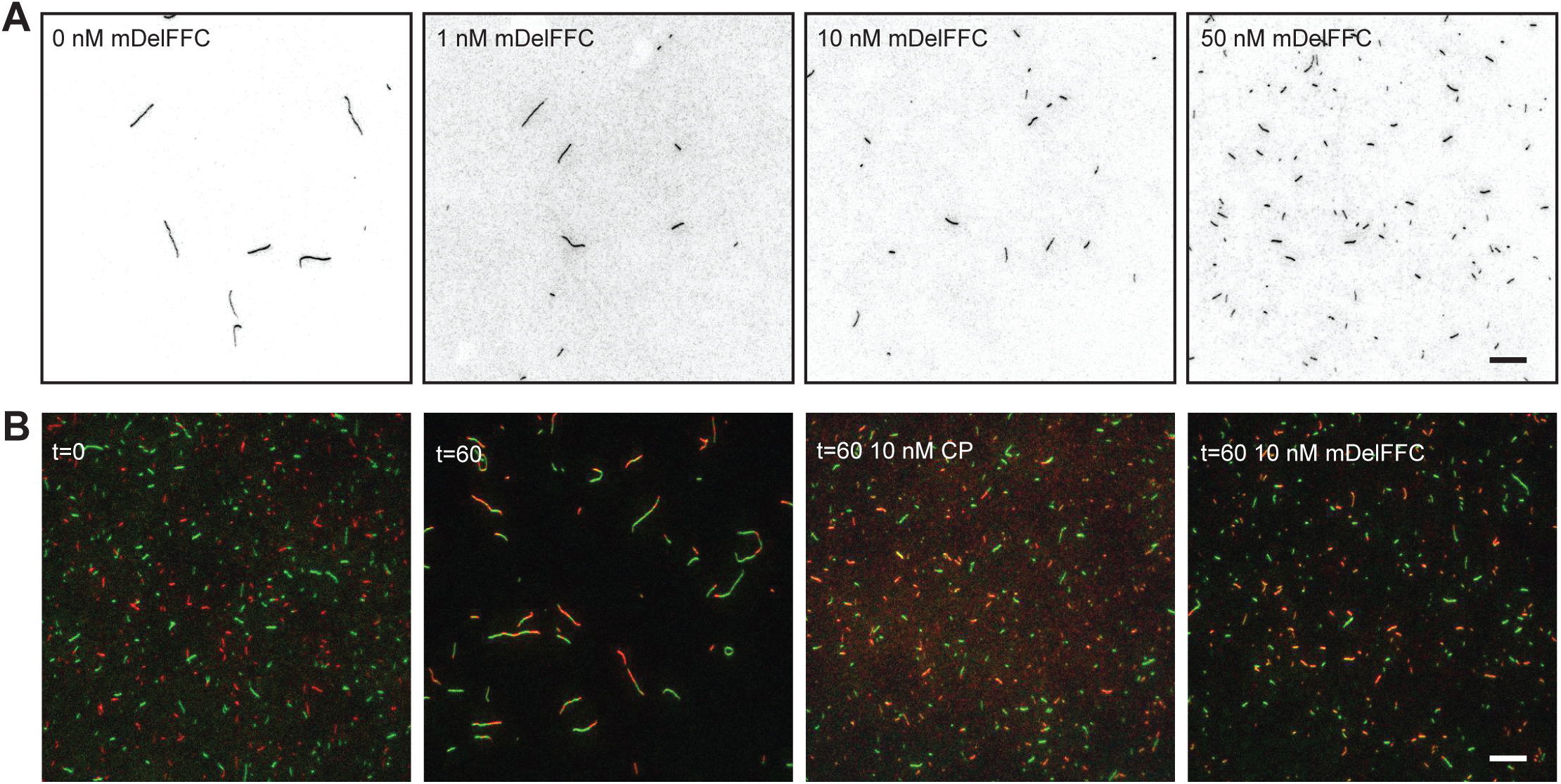
Delphilin is a nucleator. A) Actin assembly (2 μM) was triggered by addition of salts and the indicated concentration of mDelFFC. After 5 minutes, filaments were stabilized with Alexa488-phalloidin, diluted, and adsorbed to poly-L-lysine coated coverslips. Typical fields of view are shown for a range of concentrations, as indicated. Increasing concentrations of Delphilin resulted in increasing numbers of shorter filaments, indicating that Delphilin nucleates. B) A total of 0.25 μM Alexa488- and Alexa647-phalloidin-stabilized filaments were mixed and sheared. Immediately after shearing (t = 0) filaments were short and monochromatic. After an hour (t = 60), the filament length depended on the conditions. Reannealing is readily apparent when only buffer is added. Short filaments, indistinguishable from t = 0, are seen in the presence of 10 nM capping protein or Delphilin, demonstrating that Delphilin remains bound to filament ends for long periods of time. Scale bars = 10 μm

Because we observed short filaments in the nucleation assays, we hypothesized that Delphilin remained tightly associated with the barbed end. To test this hypothesis, we performed re-annealing assays over a range of mDelFFC concentrations. We mixed and sheared two populations of actin filaments stabilized with different fluorescently labeled phalloidins in the absence and presence of mDelFFC. Capping protein was used as a positive control. We obtained images immediately after mixing actin filaments (t=0) and an hour after incubation (t=60) (Figure 2B). Immediately after shearing, the overall length of filaments was short and filaments were single colors, revealing that no re-annealing events had occurred. After an hour, filaments were significantly longer and were dual labeled indicating that re-annealing of filaments occurred when no other protein was added. As expected, incubation with 10 nM CP prevented re-annealing. 10 nM mDelFFC produced results similar to 10 nM CP. These results suggest that Delphilin remains associated with barbed ends for long periods of time, similar to other formins.

### Delphilin is a permissive elongation factor

To address how Delphilin modulates the elongation rate of actin filaments, we first measured the affinity of mDelFFC for barbed ends, using a depolymerization assay. When filamentous actin was diluted to 0.1 μM in the presence of Delphilin, depolymerization was potently inhibited, reflecting tight binding (Figures 3A and S2C; K_i_ = 0.6 nM). We next used seeded elongation assays to bypass the nucleation step. We mixed 0-60 nM mDelFC or 0-350 nM hDelFFC and 0.5 μM G-actin with 0.25 μM actin seeds to monitor elongation. Both constructs tested, hDelFFC and mDelFC, potently inhibited barbed end elongation (Figures 3B and S2D). hDelFFC inhibited ~96% of elongation with a IC_50_ of 13 nM (Figure 3D). mDelFC bound barbed ends with a slightly higher affinity (IC_50_ = 7 nM) and inhibited elongation to the same degree (97%; Figure 3D). Tight barbed end binding by Delphilin and inhibition of both elongation and depolymerization are consistent with the short filaments observed in the TIRF nucleation assays and Delphilin’s ability to inhibit re-annealing (Figure 2).

**Figure 3.**
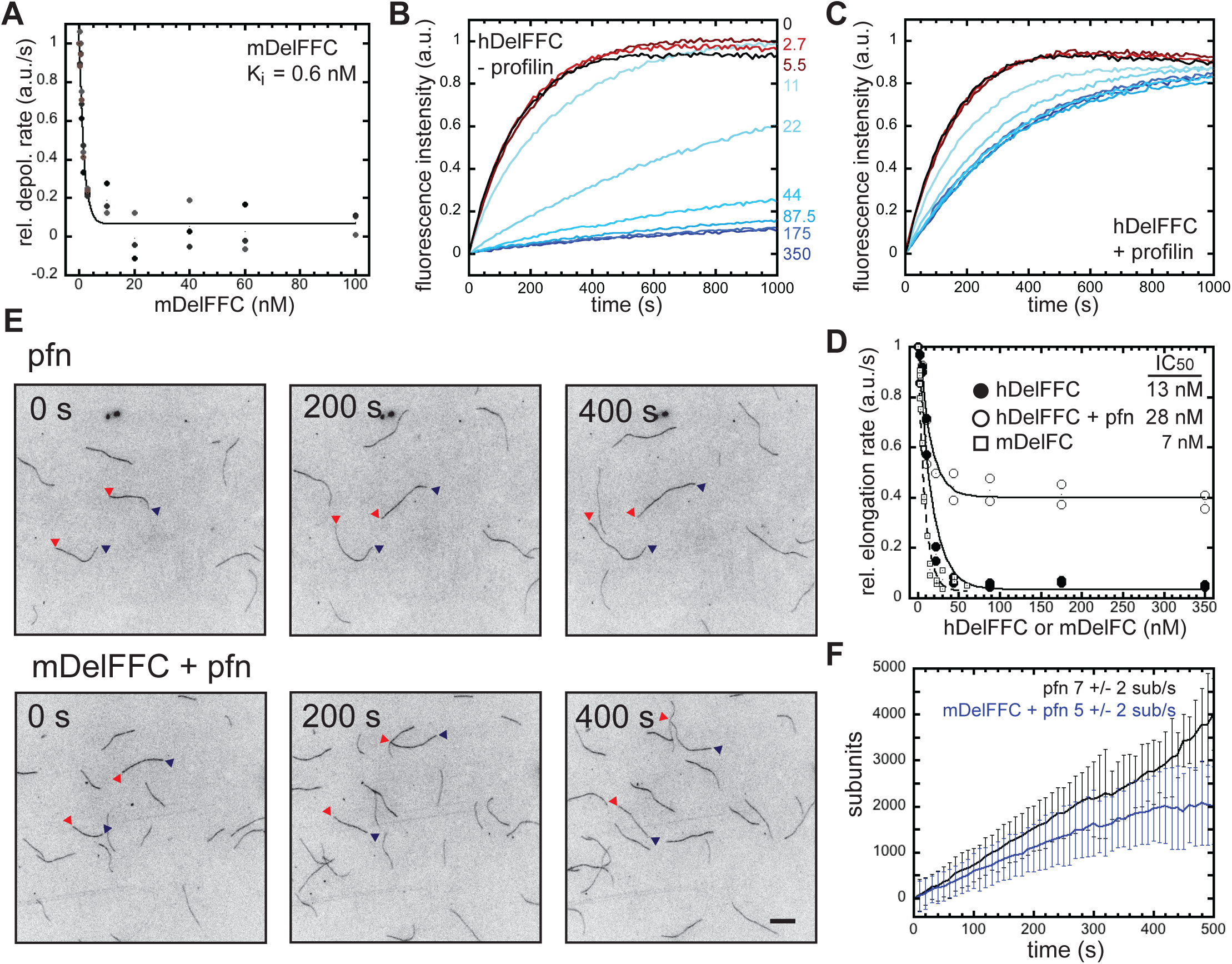
Delphilin is a permissive elongation factor. A) The initial rate of depolymerization is plotted versus the concentration of mDelFFC added. All of the data are shown. Lines are fits to averages of the data. The K_i_ reported (0.6 nM) is the average of fits to the three independent experiments. Representative raw data are shown in Figure S2C. B) Seeded elongation assays with 0.25 μM seeds, 0.5 μM actin monomers (10% pyrene labeled) and the indicated concentration of hDelFFC. Delphilin inhibits barbed end elongation. Quantification is shown in (D). C) Seeded elongation assays as in B except with 1.5 μM profilin added. D) Initial rate of elongation versus concentration of Delphilin. Data from two experiments each with or without profilin are shown. The concentration required to inhibit elongation 50% was similar in each case (13 vs. 28 nM). The degree of inhibition differed (96% vs. 64%), with hDelFFC inhibiting essentially all elongation at saturation and profilin relieving inhibiton. E) Direct observation of filament elongation by TIRF microscopy. Conditions: 2 μM actin (15% Cy3b-actin), 6 μM profilin-1, and 0.4 nM mDelFFC. Scale bar = 10 μm F) Measurement of elongation rates from TIRF experiments in the presence or absence of mDelFFC. Elongation may be slightly slower in the presence of mDelFFC but the difference is not statistically significant.

Barbed end elongation is often accelerated in the presence of formins and profilin. We therefore added profilin to the seeded elongation assay with hDelFFC (Figure 3C). In the presence of profilin, elongation was only ~64% inhibited as opposed to 96%, while the affinity for the barbed end was similar (IC_50_ = 28 nM; Figure 3D). The decreased inhibition indicated that Delphilin allows actin assembly in the presence of profilin. We used TIRF microscopy to directly monitor single-filament elongation in the presence of mDelFFC. Under these conditions (6 μM human profilin-1 and 2 μM Mg-G-actin), the average elongation rate was 7 ± 2 sub/s (Figure 3E,F). Addition of mDelFFC had little effect on the rate of filament elongation: 5 ± 2 sub/s (Figure 3E,F). These data indicate that Delphilin “permits” but does not enhance elongation in the presence of profilin, similar to what was observed for *Drosophila* Daam (Barkó *et al.*, 2010). Failure to accelerate elongation may be explained by the composition of the polyproline stretches within the FH1 domain and/or gating of the FH2 domain (see Discussion). Based on these observations, we conclude that the majority of activity observed in our original pyrene actin assembly assays (Figure 1B) reflects nucleation by Delphilin.

The reannealing data (Figure 2B) indicate that Delphilin remains bound to barbed end for a long time, like other processive formins. We next asked whether Delphilin remains associated with barbed ends under conditions where actin is elongating. To do so we used the pyrene assay and added both profilin and capping protein. As expected, addition of profilin and capping protein prevented spontaneous actin polymerization. Increasing concentrations of DelFFC were able to antagonize the inhibitory activity of profilin and capping protein and assemble actin filaments (Fig. S2E). We conclude that Delphilin remains associated with growing barbed.

### Delphilin is a weak actin bundler

The C-terminal half of multiple formins bind and/or bundle actin filaments (Michelot *et al.*, 2005; Harris *et al.*, 2006; Quinlan *et al.*, 2007; Schönichen *et al.*, 2013). We first determined how tightly mDelFFC binds the sides of actin filaments. To do so, we performed high-speed cosedimentation assays, using 0.5 μM mDelFFC and varying concentrations of phalloidin stabilized actin filaments. The data were noisy but reflected an affinity on the order of 9 +/- 4 μM (Fig. 4A). Consistent with weak side binding, at saturation only one mDelFFC dimer was bound per 20 actin monomers. Another group reported much tighter binding (Dutta *et al.*, 2017). When we reanalyzed their data, fitting it with a hyperbolic curve instead of a line, the K_d_ was ~3 μM, which is in reasonable agreement with our findings.

**Figure 4.**
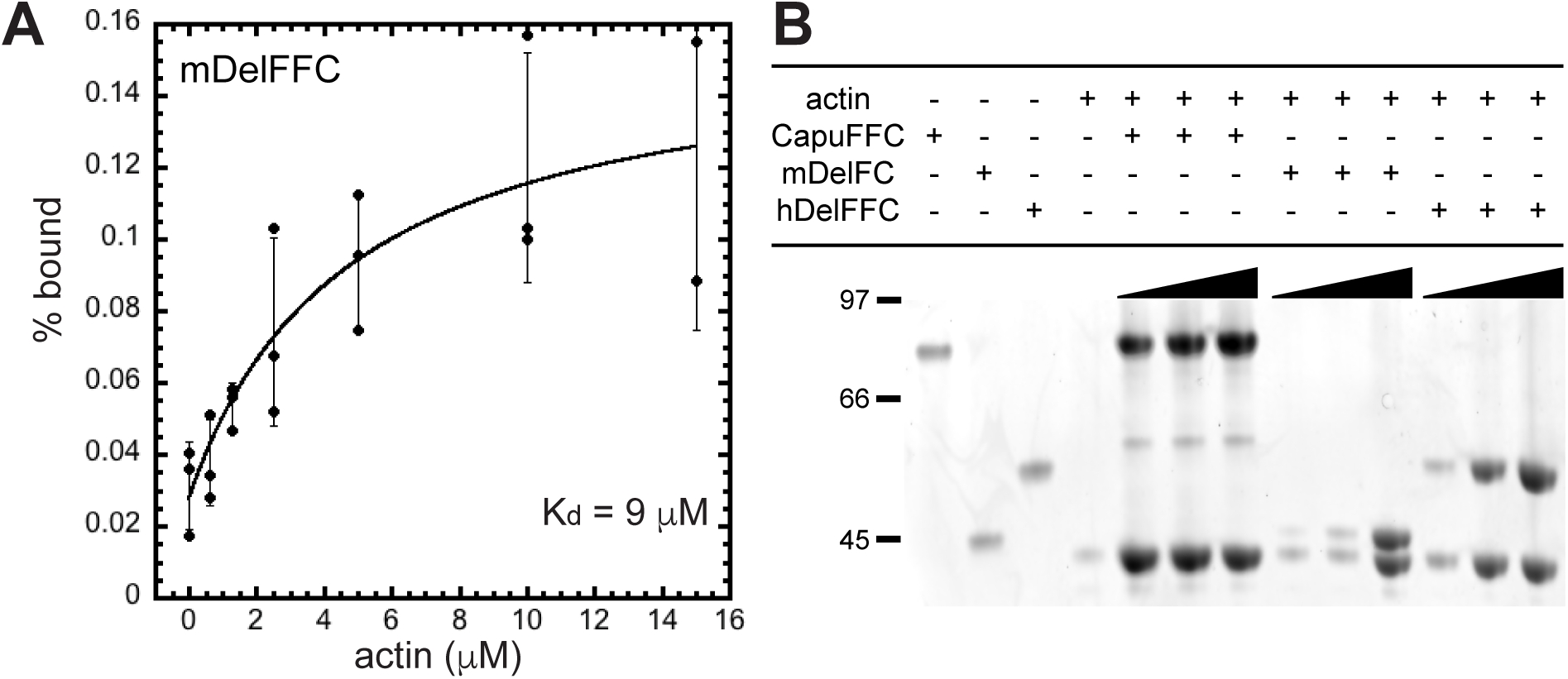
Delphilin is a weak actin bundler. A) Using cosedimentation, we measured mDelFFC binding to actin filament sides. Data from three independent experiments are shown. Lines are fits to averages of the data. The K_d_ reported (9 μM) is the average of fits to the three independent experiments. B) Low speed sedimentation assays demonstrate that mDelFC and hDelFFC bundle actin filaments. Consistent with weak side binding, bundling is weak compared to CapuFFC. Varying amounts of formin (1.25, 2.5, and 5 μM) were incubated with 5 μM F-actin for 30 min at 25°C. Bundles were sedimented by centrifugation at 12,000 ×*g* for 15 min at 4°C.

Only some formins that bind filament sides also bundle actin (Harris et al 2006). We used low-speed cosedimentation to test for bundling by mDelFC and hDelFFC (Fig. 4B). Both constructs have weak bundling activity compared to the *Drosophila* formin Cappuccino (Capu), a known filament bundler (Rosales-Nieves *et al.*, 2006; Quinlan *et al.*, 2007; Vizcarra *et al.*, 2014). While actin is found in the low-speed pellet in at all ratios of CapuFFC:F-actin (1:4, 1:2, 1:1), it is only found in the pellet for 1:1 Delphilin:F-actin. We interpret this data as weak actin bundling activity of Delphilins.

### Delphilin lacks a functional tail

The tails of several formins, including mDia1, mDia2, Bni1, Bnr1, FRL, and Capu, are important for nucleation and processivity in addition to autoinhibition (Gould *et al.*, 2011; Vizcarra *et al.*, 2011; Heimsath and Higgs, 2012). Tail lengths vary a great deal among mammalian formins (Figure 5A). Based on alignments and our homology model, Delphilin does not have a C-terminal tail (Figure 5A,B). In fact, the predicted last α-helix in the FH2 domain of Delphilin, αT, is shorter than αT of other formins that have been crystalized. In addition, only eight residues remain beyond the predicted helix (Figure 5B,C). To further investigate the Delphilin tail, or lack thereof, we compared mDelFC to constructs lacking 5 to 25 residues. mDelFCΔ25 was insoluble but the rest of the constructs were readily purified with the same protocol (Figure S1A). We first assessed the impact of the truncations by measuring thermal stability of the proteins. The constructs were essentially indistinguishable until 20 residues were removed, a truncation that is predicted to be well into the αT helix (Figure 5B,C). Wild type mDelFC has a melting temperature of 46.5°C and mDelFCΔ20 melts at 40.5°C (Figure 5D). In pyrene-actin assembly assays, mDelFC activity was not affected by truncation (Figure 5E). In addition, barbed end binding was unaffected; mDelFC and mDelFCΔ20 inhibited seeded elongation to the same extent (Figure 5F). Thus, Delphilin does not have a tail that contributes to actin nucleation or end binding.

**Figure 5.**
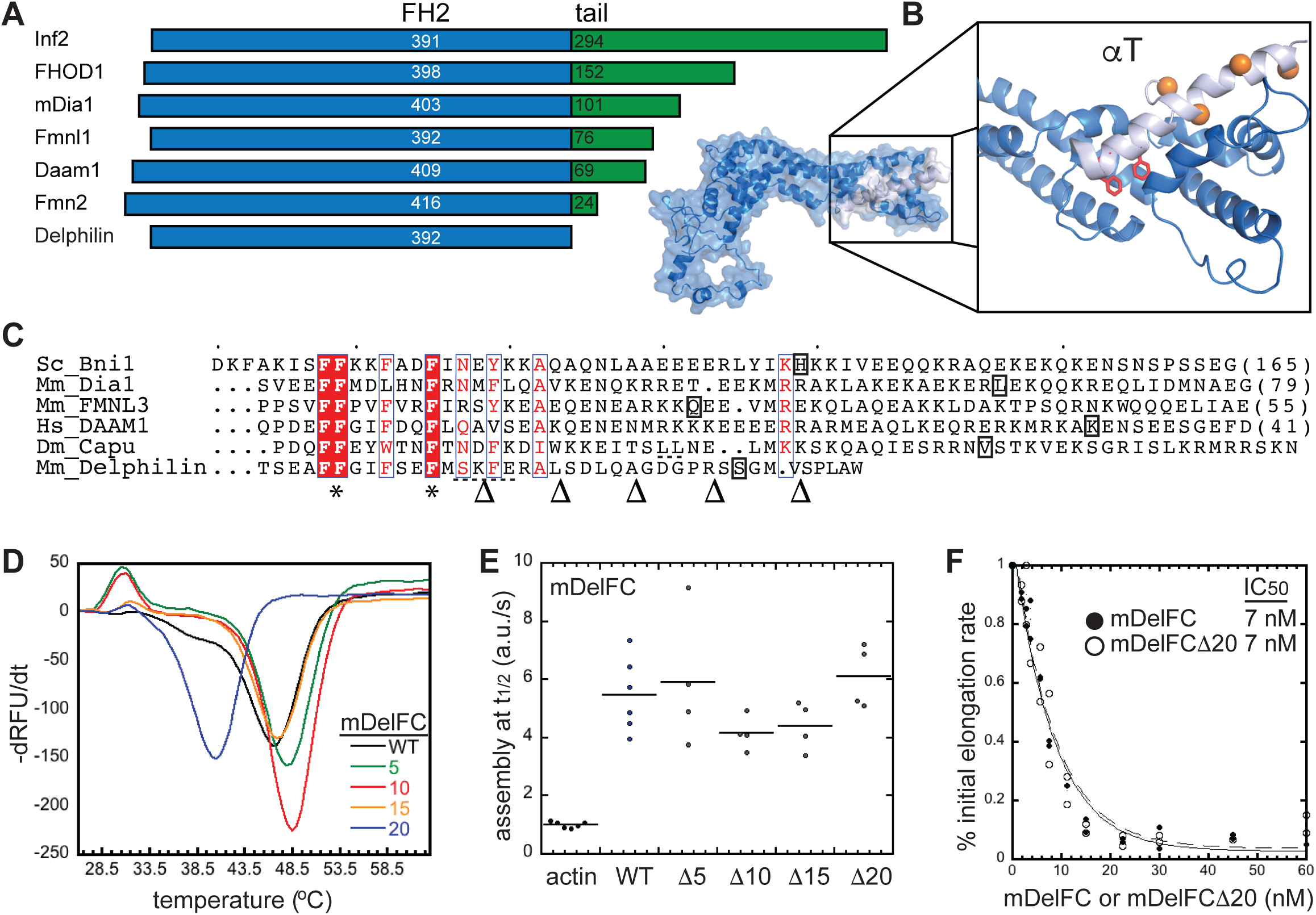
Delphilin lacks a functional tail. A) Comparison of tail length among the seven classes of mammalian formins. The blue FH2 domains are aligned at their C-termini. The tails are in green. Numbers indicate the length of each domain. B) Homology model of Delphilin-FH2 based on the Daam1 crystal structure (2J1D). The C-terminus of the structure is expanded to show the short final helix (αT, light blue). Delphilin’s final eight residues are absent from the model. Conserved phenylalanines at the base of αT are shown as red sticks. Orange residues (balls) indicate where the Δ10 - Δ25 truncations were made. (Δ5 is not in the structure.) C) Alignment of αT sequences based on crystal structures (Bni1 (1UX5), mDia1 (1V9D), FMNL3 (4EAH), DAAM1 (2J1D)) and homology models (Capu (Yoo *et al.*, 2015), Delphilin). * indicates phenylalanines conserved in all formins which are indicated in red in (B). The boxed letter is the last residue observed for each structure. Dashed underlines indicate a predicted non-helical insert in the middle of αT. Arrowheads at the bottom indicate where truncations were made for experiments in (D-F) D) Thermal stability of mDelFC truncations compared to wild type (the number of residues removed is given in the legend). 1 μM mDelFC was mixed with Sypro Orange according to the manufacturers specifications and heated from 4°C to 95°C. Only mDelFCΔ20 was less stable than wild type. D) Assembly rates from pyrene assays with 4 μM actin and 150 nM mDelFC construct, as indicated. The rates did not differ significantly for any of the constructs, including mDelFCΔ20. E) Inhibition of seeded elongation by mDelFC and mDelFCΔ20. The absence of the last 20 residues has no effect on mDelFC’s ability to inhibit barbed end elongation (IC_50_ = 7 nM for both constructs). Data from two independent repeats are shown. Representative raw data are shown in Figure S2D.

### Delphilin is not autoinhibited

Autoinhibition is a hallmark of formins and occurs through conserved DID and DAD domains (Li and Higgs, 2005; Goode and Eck, 2007). Fmn-family formins lack canonical DID and DAD domains but are auto-inhibited (Bor *et al.*, 2012). Delphilin is one of two other formin families lacking these domains. As described above, Delphilin lacks a tail, which is the typical site of the DAD domain. Despite this, we asked whether Delphilin can be autoinhibited. To do so, we purified the N-terminal half of α-Delphilin (α-mDelNT). Circular dichroism revealed the presence of a large amount of secondary structure, α-helix and β-sheet (>50%) suggesting that the α-mDelNT construct is folded (Figure S1C). Addition of α-mDelNT to actin alone does not affect actin polymerization. Titrating α-mDelNT with mDelFFC revealed no change in the polymerization activity of mDelFFC (Figure 6A), indicating that actin assembly is not inhibited by the intramolecular interaction typical of formins. In order to assess whether there was an interaction between the two halves of the protein, we used size exclusion chromatography. When mixed together, α-mDelNT and mDelFC migrate independently through the column (Figure 6B).

**Figure 6.**
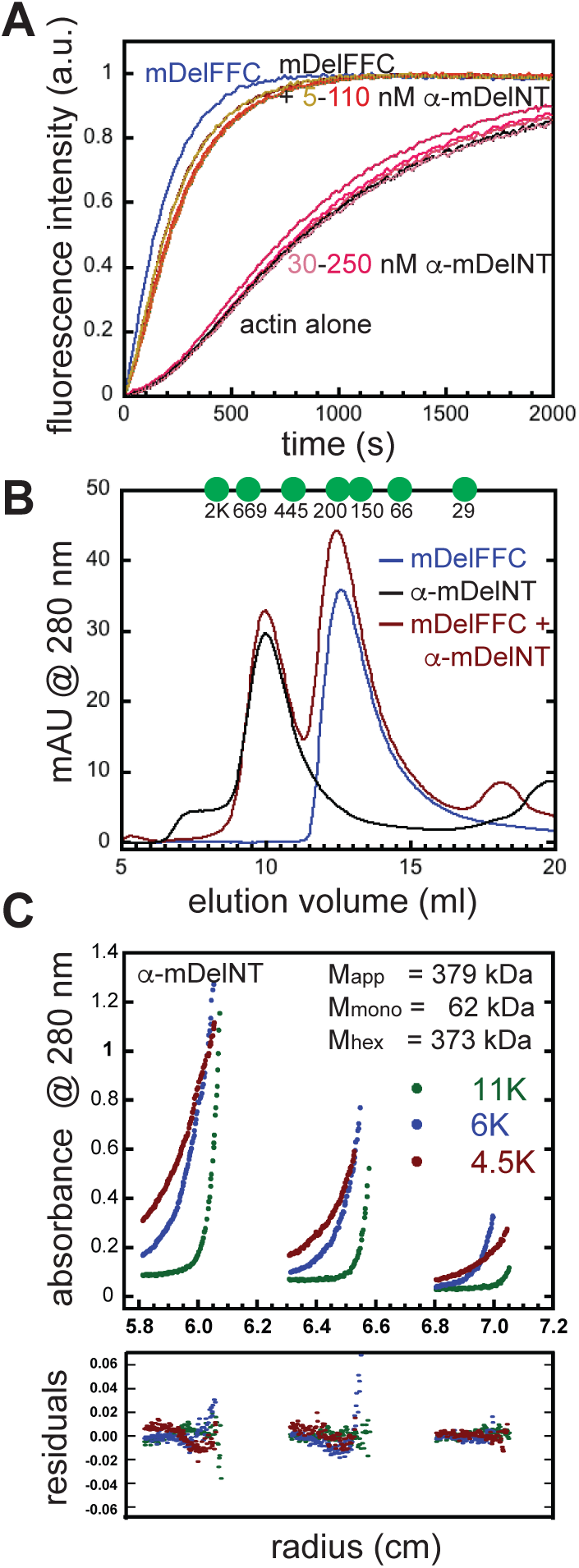
Delphilin is not autoinhibited. A) The N-terminal half of α-Delphilin (mDelNT) has no effect on actin assembly whether or not mDelFFC is present. Conditions: 4 μM actin (10% pyrene labeled), 30 nM mDelFFC, and a range of α-mDelNT concentrations, as indicated. B) Size exclusion chromatography was performed on mDelFC (blue), α-mDelNT (black) or both (red) with a Superdex S200 30/100 GL column. No differences were detected when the two proteins were mixed versus alone. Elution peaks of standards are indicated with green circles. Y-axis is milli-absorbance units (mAU) measured at 280 nm. C) Equilibrium sedimentation data for three concentrations of α-mDelNT at three speeds were globally fit with a single species model. The data indicate that α-mDelNT is a hexamer. Values for the apparent mass (M_app_) and the masses predicted for α-mDelNT monomers (M_mono_) and hexamers (M_hex_) are given.

The N-terminal halves of most formins are dimers (Li and Higgs, 2005; Rose *et al.*, 2005; Bor *et al.*, 2012). To determine the molecular weight of α-mDelNT, we first used size exclusion chromatography. α-mDelNT eluted as a single peak at a volume that predicts multimer of 7 +/ 1 subunits (Figure 6B). Because size exclusion chromatography is sensitive to shape as well as size, we turned to equilibrium sedimentation. Using single species modeling we calculated the molecular weight of α-mDelNT to be 379.5 kDa. The predicted molecular mass of a hexamer is 373.5 kDa, a 2% deviation from our measurement (Figure 5C). The fit was not improved by a monomer-hexamer model. We note a trend in the regressions. This could reflect one or both of the following: 1) there is a contaminant that is smaller than an α-mDelNT; 2) the hexamer is not homogenous. In both cases, the data support the presence of a dominant species as opposed to variable aggregates of unfolded protein. Thus, Delphilin may be exceptional among formins in oligomer state as well.

## Discussion

### Actin assembly by Delphilin

While most formins slow actin elongation in the absence of profilin, only Cdc12, *Drosophila* Daam (dDaam), and now Delphilin completely cap barbed ends (Kovar *et al.*, 2003; Barkó *et al.*, 2010). In the presence of profilin, most formins accelerate elongation. This is true for Cdc12, which accelerates elongation about 50% over actin alone (Kovar *et al.*, 2006). In contrast, inhibition of elongation is relieved but elongation is not accelerated by Delphilin or Daam, placing them at one end of a spectrum of possible actin elongation activities ((Barkó *et al.*, 2010); and this study). Phylogenetic analysis of formins based on FH2 domains, places Delphilin closer to Diaphanous, one of the most potent elongation factors, than to Daam (Pruyne, 2016). Thus the profilin-binding FH1 domain, and not the FH2 domain, is likely to be largely responsible for this extreme activity. The number and lengths of polyproline stretches and their distances from the FH2 domain all contribute to activity levels (Paul and Pollard, 2008; Courtemanche and Pollard, 2012). For example, experiments and simulations show that higher affinity proline stretches spaced 18 residues from the FH2 domain accelerate elongation less effectively than lower affinity proline stretches with the same spacing (Courtemanche and Pollard, 2012). In fact, the closest polyproline stretch is usually a relatively weak profilin binder, presumably facilitating actin handoff to the FH2 domain (Courtemanche and Pollard, 2012). Among the mammalian formins, the number of polyproline stretches range from 2 to 19, in Delphilin and Fmn2, respectively. The linker regions, the distance between the last polyproline repeat and the beginning of the FH2 domain vary from 10 to 47, in FHOD1 and Fmn2, respectively. The linkers of both dDaam and mDelphilin are short, 16 and 11 residues, respectively. In contrast, the Cdc12 linker is 26 residues. Even in their fully-extended conformations, the short linkers of dDaam and mDelphilin are equivalent to or shorter than the diameter of an actin monomer, which could hinder handoff. Further, Delphilin has only two polyproline stretches and the longer of the two is proximal to the start of the FH2 domain, which could also hinder handoff. Thus Delphilin is not well designed to enhance elongation, which is consistent with our observations.

We also note the decrease of activity in mDelFC relative to mDelFFC. Nucleation activity is generally attributed to the FH2 domain. In fact, while the FH2 domain is sufficient, nucleation may be influenced by the FH1 domain in some cases. For example, comparison of Bni1-FC and FFC constructs reveals a large difference in nucleation strengths (Pruyne *et al.*, 2002; Moseley *et al.*, 2004). In contrast, the activity of Capu-FC and -FFC constructs are indistinguishable (Quinlan *et al.*, 2007). However, the Capu-FC construct is very unstable compared to the -FFC construct (our unpublished observations). It may be that the FH1 domain does not directly contribute to nucleation but instead influences FH2 structure and, therefore, activity. The contribution of the FH1 domain to nucleation is interesting in light of the fact that several formins, including Delphilin, are only active elongation factors in the presence of their FH1 domains and profilin (Kovar *et al.*, 2003; Michelot *et al.*, 2005; Barkó *et al.*, 2010). It is difficult to distinguish whether the FH1 domain strictly contributes by delivering profilin-actin to the growing filament or if the profilin-bound FH1 domains alter gating, for example. Given Delphilin’s unusual FH1 domain and lack of a tail, its FH2 domain could be an interesting base around which to further investigate how FH1 and tail domains enhance actin assembly.

Another group recently reported a biochemical study of Delphilin (Dutta *et al.*, 2017). They described Delphilin as a barbed end capping protein, which is consistent with our data, in that Delphilin tightly binds barbed ends and, under nucleating conditions, only pointed end growth is observed in the absence of profilin. They also report that nucleation is completely inhibited in the presence of profilin. We see strong inhibition of nucleation by profilin as well but detect activity. They observed some relief of inhibition in seeded elongation assays plus profilin but did not examine elongation directly. We believe the actin isoform used is the major source of the differences, given the very weak activity of Delphilin with rabbit skeletal actin.

### Regulation of Delphilin

Unlike, all other characterized formins, Delphilin does not appear to be autoinhibited. The absence of an appreciable tail-domain C-terminal to the FH2 domain eliminates the possibility of canonical DID/DAD autoinhibition (Li and Higgs, 2005; Goode and Eck, 2007). However, an intramolecular interaction involving the FH2 domain could also inhibit actin assembly. We detected no evidence of such an interaction.

The N-terminal halves of mDia1, Capu, and most other formins are dimers (Otomo *et al.*, 2005; Rose *et al.*, 2005; Bor *et al.*, 2012). In contrast, the N-terminus of Delphilin forms a hexamer, further distinguishing it from canonical DID/DAD regulation. Multimerization is reported for other PDZ domain containing proteins. Specifically, PSD-95 has been shown to multimerize, clustering other proteins at membranes (Kim *et al.*, 1995). Two N-terminal cysteines (Cys-3 and Cys-5) in the PDZ domain of PSD-95 are required for multimerization (Hsueh and Sheng, 1997). There are two cysteines in the N-terminal portion of Delphilin-PDZ (Cys-3 and Cys-29) suggesting the multimerization may be mediated by a mechanism similar to that of PSD-95. While our data, circular dichroism, SEC, and equilibrium sedimentation, all provide evidence that α-mDelNT is folded, we cannot exclude the possibility that the purified α-mDelNT construct is partially unfolded in a manner that impacts autoinhibitory interactions.

We note that three splice variants of Delphilin have been reported, α-, β-, and L-Delphilin (Yamashita *et al.*, 2005; Matsuda *et al.*, 2006). They vary in their N-terminal halves but not their C-terminal halves, meaning that none has a C-terminal “tail”. The α variant is the shortest transcript and can be palmitoylated at Cys-3 (Matsuda *et al.*, 2006). The first five residues of α- Delphilin are spliced out and replaced either with 12 residues (β-Delphilin) or 184 residues including a second PDZ domain (L-Delphilin). Given the small difference between α- and β- Delphilin, we do not expect this variant to be autoinhibited. We cannot rule out intramolecular interactions in the case of L-Delphilin. Immunofluorescence images in cultured neurons indicate that the N-termini are important for localization. α-Delphilin is specifically localized in dendritic spines, whereas β-Delphilin is relatively diffuse in both dendritic spines and the shafts of spines (Yamashita *et al.*, 2005; Matsuda *et al.*, 2006). In contrast, L-Delphilin forms clusters that are predominantly within the neuron shafts (Matsuda *et al.*, 2006). Coordination of localization and activity has been reported for other formins (Seth *et al.*, 2006; Ramalingam *et al.*, 2010). Whether this is the case for Delphilin remains to be determined. In the absence of autoinhibition, one can only speculate about regulation. Two likely possibilities are interactions with an unknown partner and/or post-translational modification. Although, Delphilin is a weak elongation factor, it is a relatively strong nucleator, making it unlikely that its activity goes unchecked.

Two new families of formins were recently identified, Multiple Wing Hairs-relate Formin (MWHF) and Pleckstrin Homology domain-Containing Formins (PHCF) (Pruyne, 2016). Formins in the MWHF family have conserved DID and DAD domains and are likely to be autoinhibited. Formins in the PHCF family do not have these domains. Instead, they have Pleckstrin homology domains. Most have more than one PH domain and some have C-terminal PH domains in addition to or instead of N-terminal PH domains. It is interesting to note that PDZ and PH domains both play roles in coordinating proteins at the membrane. Perhaps the Delphilin and PHCF families of formins evolved, function, and/or are regulated in similar ways, though they are not closely related based on FH2 sequences.

### Role in dendritic spines

How the actin cytoskeleton is regulated within dendritic spines affects their structure and function (Hlushchenko *et al.*, 2016; Lei *et al.*, 2016). We set out to determine the biochemical characteristics of the formin, Delphilin, because it is the only formin specifically expressed in dendritic spines. mDia2, Daam1, and the Arp2/3 complex are all implicated in distinct stages of spine formation, including filopodia formation and head expansion (Salomon *et al.*, 2008; Hotulainen *et al.*, 2009). Interestingly, Delphilin is not required for spine formation (Takeuchi *et al.*, 2008). Instead it binds to the C-terminus of GluRδ2, a receptor crucial to LTD (Miyagi *et al.*, 2002). Other PDZ domain proteins bind the same site of GluRδ2 and have been implicated in transduction of signals leading to LTD (Kohda *et al.*, 2007). In contrast, induction of LTD is enhanced in the absence of Delphilin, suggesting that it plays an inhibitory role in this process (Takeuchi *et al.*, 2008). Competition for GluRδ2-binding is an obvious possibility. At this time, we can only speculate about mechanism and the significance of nucleation by Delphilin. For example, Delphilin could build a structure that stabilizes its interaction with GluRδ2, limiting the downstream response to activation of this receptor. Based on the work presented here, the knockout mouse already made by others, and tools available to perform “rescue” experiments in cerebellar slices, it is now possible to directly test whether nucleation activity is essential (Kohda *et al.*, 2007; Takeuchi *et al.*, 2008).

Finally, we consider the other formin known to be a poor elongation factor, dDaam (Barkó *et al.*, 2010). It is implicated in a range of roles, including tracheal tube formation, growth cone filopodia formation, sarcomerogenesis, axonal growth, and even non-canonical Wnt signaling (Sato *et al.*, 2006; Salomon *et al.*, 2008; Barkó *et al.*, 2010; Higashi *et al.*, 2010; Li *et al.*, 2011). dDaam could be specialized to build small structures needed in a wide number of cells and processes. Alternatively, elongation by formins like dDaam and Delphilin may be enhanced by additional binding partners in a manner similar to that recently described for mDia1 and CLIP-170 (Henty-Ridilla *et al.*, 2016).

## Methods

### DNA construct

mDelFFC (aa 559-1024; NP_579933.1), mDelFC (aa 626-1024), and α-mDelNT (aa 1-558) were generated by PCR amplification off a mDelphilin-FL template which was kindly provided by S. Kawamoto (Chiba University). These constructs were subcloned into a slightly modified pET-15b+ plasmid with a BamHI site immediately following the N-terminal His tag. mDelFC point mutations, I718A and K868A, as well as the mDelFC tail truncations were introduced by QuikChange Site Directed Mutagenesis (Stratagene, Santa Clara, CA). hDelFFC (aa: 744-1211; NP_01138590) was assembled from synthetic DNA, which was codon optimized for E. coli expression (IDT), and cloned into pET-15b+.

### Protein expression and purification

All Delphilin constructs were expressed in Rosetta- or Rosetta- II (DE3) pLysS-competent cells. Cells were grown in Terrific Broth at 37°C until they reached an O.D. 600 nm of 0.7-1.0, the temperature was then lowered to 18°C for ~1 h. Cells were induced with 0.25 mM isopropyl-β- D-thiogalactoside and harvested after 14-17 h. Cell pellets were then flash frozen in liquid nitrogen and stored at −80°C.

All C-terminal constructs were purified and stored using published protocols, with the exception that mDelFFC was stored in 10 mM Tris, pH 8, 50 mM KCl, 1 mM DTT and 50% glycerol (Vizcarra *et al.*, 2011). Briefly, protein was purified on a Talon column followed by a monoQ column. α-mDelNT cell pellets were resuspended in lysis buffer (50 mM Tris, pH 7, 150 mM KCl, 1mM dithiothreitol [DTT]) supplemented with 2 mM phenylmethanesulfonyl fluoride (PMSF) and 1 μg/mL DNaseI. All following steps were carried out on ice or at 4°C. Resuspended cells were passed three times through a microfluidizer (Microfluidics, Newton, MA). The lysate was centrifuged at 30,000 x *g* for 25 minutes. Supernatant was then added to 1 mL of Talon resin (GE Healthcare) for 1 h. α-mDelNT was eluted from resin with 200 mM imidazole and spin concentrated in an Amicon 10-kDa-molecular weight cut off centrifugal filter unit and gel filtered on a Superdex 200 10/300 GL column (GE Healthcare) in 10 mM Tris, pH 8, 50 mM KCl, and 1 mM DTT. Fractions of α-mDelNT were pooled based on SDS-PAGE analysis and then dialyzed into 10 mM Tris, pH 8, 50 mM KCl, 1 mM DTT and 50% glycerol. Aliquots were flash frozen in liquid nitrogen and stored at −80°C. The resulting samples were >95% pure (Supplemental Figure 1A).

Protein concentrations of C-terminal constructs are reported in terms of dimer concentration α- mDelNT concentration is reported as a monomer. Protein concentrations were determined by analyzing five serial dilutions of the sample and a standard by SDS-PAGE. The gels were stained with SyproRed (Invitrogen) and imaged on a Pharos FX Plus molecular imager with Quantity One software (Bio-Rad).

*Acanthamoeba castellani* actin purification and labeling with Oregon green 488 and Cy3b was performed according to previously published protocols (MacLean-Fletcher and Pollard, 1980; Bor *et al.*, 2012). Rabbit skeletal actin was a generous gift from the Reisler lab (UCLA). Cytoplasmic actin purified from human platelets (85% beta, 15% gamma) was purchased from Cytoskeleton, Inc. (APHL99-A). Unless otherwise indicated, assays were performed with *Acanthamoeba* actin.

### Pyrene-actin polymerization assays

Pyrene-actin assembly assays were performed principally as described in (Zalevsky *et al.*, 2001). Briefly, 4 μM *A. castellani* actin (10% pyrene labeled) was incubated for 2 min at 25°C with ME buffer (final concentration, 200 μM ethylene glycol tetraacetic acid [EGTA] and 50 μM MgCl_2_) to convert Ca-G-actin to Mg-G-actin. Polymerization was initiated by adding KMEH polymerization buffer (final concentration, 10 mM Na-HEPES, pH 7.0, 1 mM EGTA, 50 mM KCl, 1 mM MgCl_2_) to the Mg-G-actin. Additional components, such as mDelFFC, hDelFFC, m-αDelNT, mDelFC and its associated mutants were combined in the polymerization buffer prior to addition to Mg-G-actin. For experiments that included profilin (*S. pombe*) a 3:1 molar ratio of profilin was added to actin before ME. For seeded elongation assays 5 μM actin was polymerized overnight. 0.5 μM *A. castellani* actin (10% pyrene labeled) and 0.25 μM F-actin seeds were used. Delphilin constructs were added to either actin alone or 1.5 μM profilin to 0.5 μM actin.

Pyrene actin fluorescence was monitored using a TECAN F200 with λ_excitation_ = 365 nm and λ_emission_ = 407 nm.

### TIRF microscopy assays

Coverslips were PEGylated by two different methods: 1) incubation with poly-L-lysine PEG or 2) treatment with 3-aminopropyltrethoxysilane followed by PEG-NHS (3% biotin-PEG-NHS) as described (Bor *et al.*, 2012). No difference in rate was observed between the two immobilization methods. Flow cells were blocked for 2 minutes with Pluronic F-127 (Sigma) and 50 μg/ml κ- casein in PBS and then washed with TIRF buffer (final concentration, 10 mM Na-HEPES, pH 7.0, 1 mM EGTA, 50 mM KCl, 1 mM MgCl_2_). Experiments were performed with 2 μM Mg-G-actin (15% Cy3b labeled), 6 μM human profilin-1, 5 nM phalloidin stabilized actin seeds, and mDelFFC in TIRF buffer, supplemented with 0.5% methylcellulose, 50 mM DTT, 0.2 mM ATP, 50 μg/ml catalase, 50 μg/ml κ-casein, and 250 μg/ml glucose oxidase, and 20 mM glucose. Images were collected every 10 s on a DMI6000 TIRF microscope (Leica). Data were analyzed using the JFilament plugin (Smith *et al.*, 2010) to Fiji (Schindelin *et al.*, 2012).

### Actin Filament Cosedimentation Assays

Binding: Actin (10 μM) was polymerized in 1× KMEH for 1 h at room temperature before adding a 1:1 molar ratio of phalloidin. mDelFFC was precleared by centrifugation at 117,000 × *g* for 20 min at 4 °C. mDelFFC (0, 0.625, 1.25, 2.5, 5, 10, and 15 μM) was incubated for 30 min at 25°C with 5 μM actin. The polymerized filaments were transferred using cut pipette tips to avoid shearing. Samples were centrifuged at 89,000 × *g* for 20 min at 4 °C. The supernatants and pellets were analyzed by SDS-PAGE. The gels were stained with SyproRed and imaged using a Pharos FX Plus molecular imager with Quantity One software (Bio-Rad).

Bundling: Each formin (1.25, 2.5 and 5 μM) was incubated for 30 min at 25°C with 5 μM (rabbit skeletal muscle) actin. The *Drosophila* formin CapuFFC was used a positive control for bundling. Samples were centrifuged at 12,000 x *g* for 15 min at 4°C. Supernatants and pellets were separated, and pellet fractions were concentrated four-fold for SDS-PAGE analysis. Protein bands were stained with SyproRed and imaged on a Typhoon FLA 7000 (GE).

### Circular dichroism

α-mDelNT was dialyzed into 10 mM Tris, 50 mM KCl, and 1 mM DTT (pH 8). Circular dichroism (CD) spectra were measured on a J-715 spectropolarimeter (Jasco, Tokyo, Japan) by averaging two wavelength scans from 195 to 280 nm. Data were analyzed using Jasco and Sofsec1 software packages, which compares the acquired data to a database of spectra from proteins with known secondary structure (Sreerama and Woody, 1993).

hDelFFC was brought to 1 μM concentration with 10 mM NaCl, 50 mM sodium phosphate, and 1 mM DTT (pH 7). CD spectra were collected on a 62DS spectropolarimeter (Aviv Biomedical, Inc.). For thermal denaturation, temperature was increased in increments of 1 °C and held at each temperature for 1 min.

### Analytical ultracentrifugation

Sedimentation equilibrium analytical ultracentrifugation was performed on α-mDelNT dialyzed in 10 mM Tris, 50 mM KCl, and 0.5 mM TCEP (pH 8.0). Three different concentrations of protein (4.8 μM, 2.5 μM, and 1.1 μM) were spun at 4°C in an Optima XL-A Analytical Ultracentrifugation system (Beckman Coulter, Brea, CA) at three speeds 4,500 rpm, 6,000 rpm, and 11,000 rpm. Scans between 24 and 28 hours were collected and analyzed.

## Acknowledgements

The authors would like to thank Dr. Susumu Kawamoto (Chiba University) for the original mouse Delphilin constructs and the Reisler lab for rabbit skeletal muscle. Thanks also to the Reisler, Quinlan, and Vizcarra labs for helpful discussions throughout the project. This work was supported by the generous support of Barnard College, the Sally Chapman Fund, and the Sherman Fairchild Foundation (CLV, GCN) and an NIGMS grant (R01 GM096133) to MEQ.

## Supplementary Figure Legends

**Figure S1.**
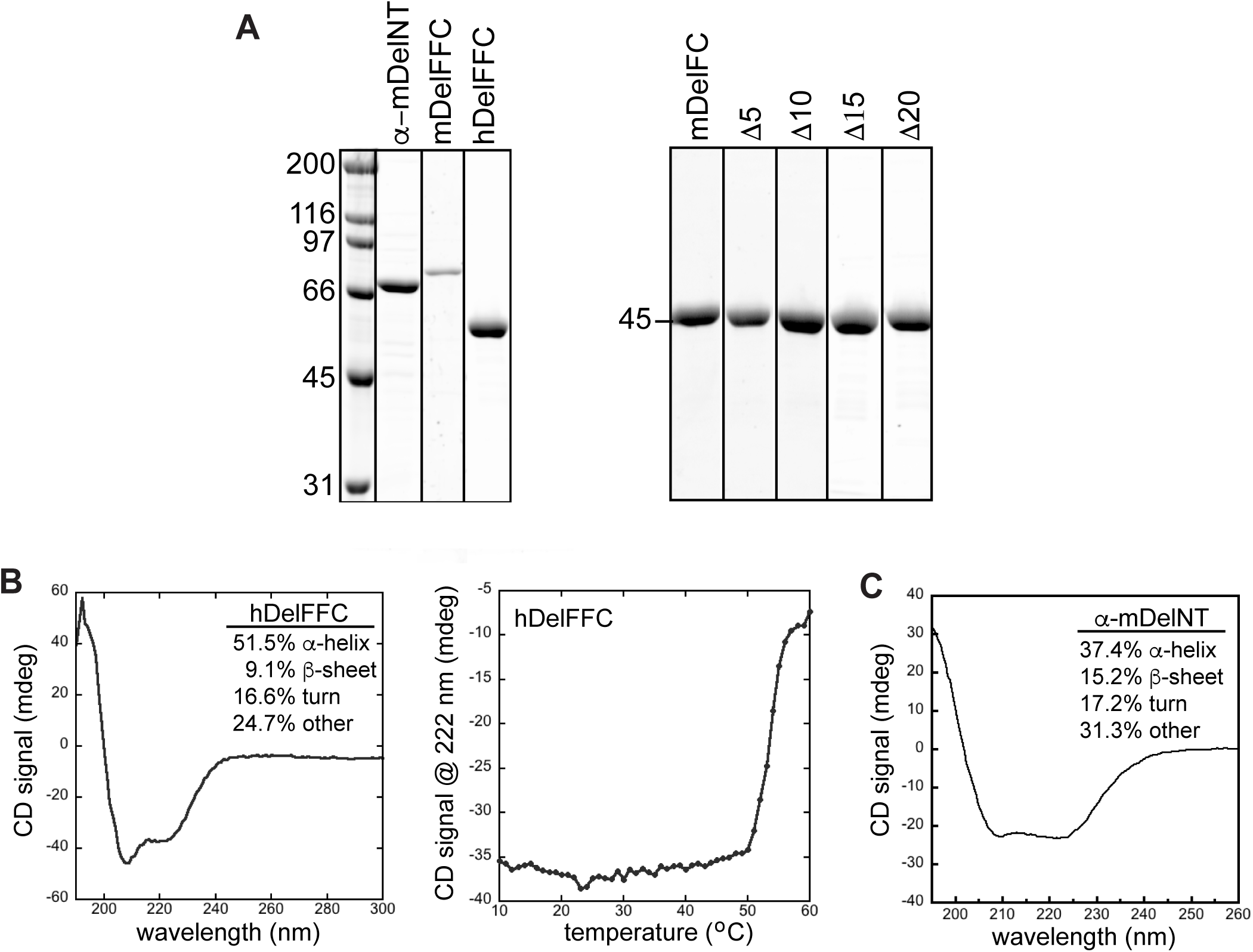
Proteins used in this study. A) SDS-PAGE gels of the constructs used in this paper: α- mDelNT, mDelFFC, and hDelFFC are grouped together. mDelFC, and truncations thereof are grouped separately. B) Circular dichroism analysis of secondary structure of 0.06 mg/ml hDelFFC. Loss of structure as a function of temperature demonstrates that hDelFFC is relatively thermal stable compared to other formins. The T_m_ (~55 °C) is about 5° higher than mDelFC, 14° higher than mDia1-FH2 (Kupi *et al.*, 2013), and 20° higher than CapuFFC (Vizcarra *et al.*, 2014). C) Circular dichroism analysis of secondary structure of 0.2 mg/ml α-mDelNT. Most of the secondary structure detected is α-helical, consistent with a PDZ domain. More than 50% of the construct has secondary structure, suggesting that it is folded.

**Figure S2.**
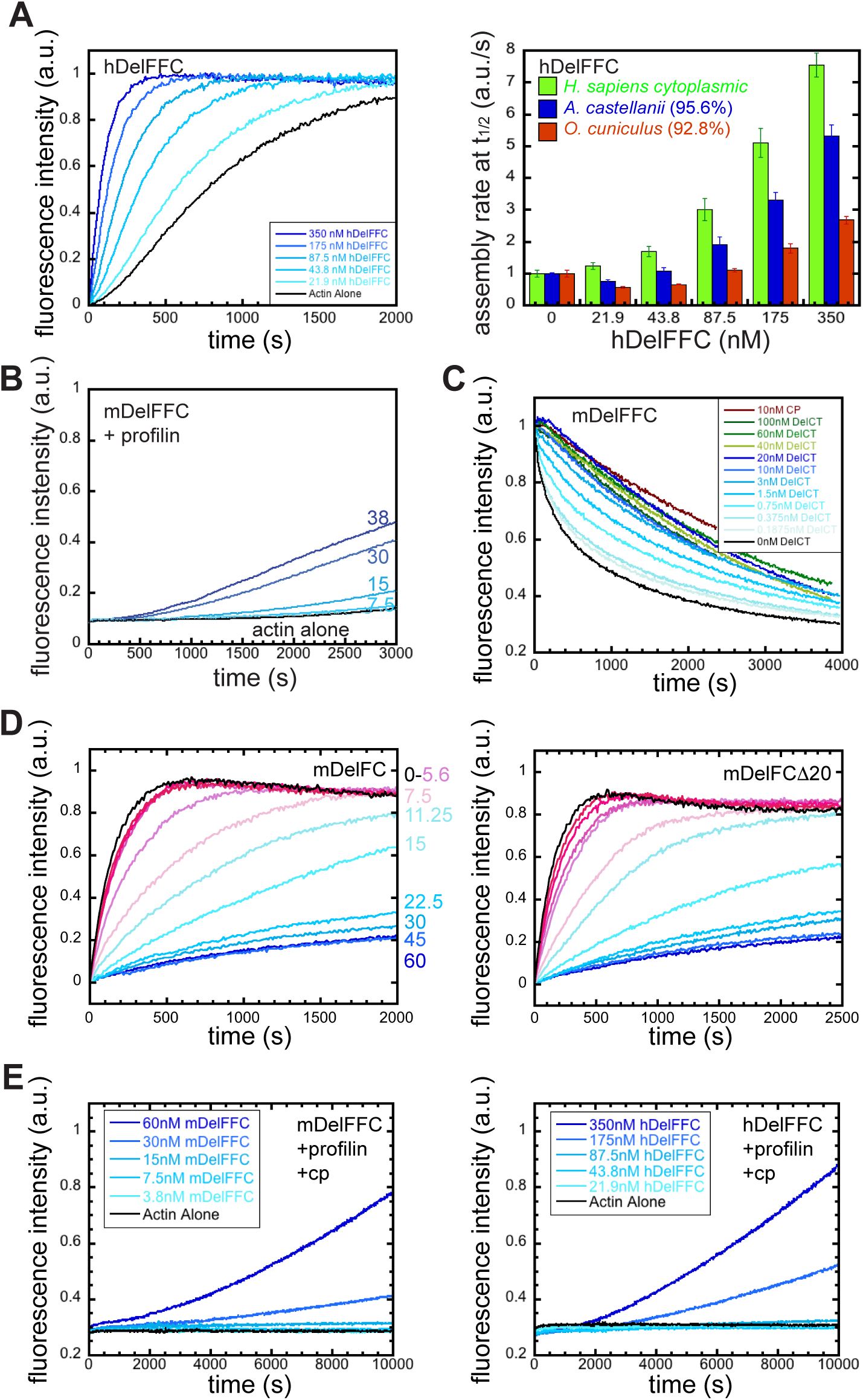
Actin polymerization and depolymerization in the presence of Delphilin. A) Left: Pyrene actin assembly assay with hDelFFC as indicated and 4 μM amoeba actin. Right: comparison of hDelFFC-stimulated assembly rates in the presence of different actin isoforms. B) Pyrene actin assembly assay with mDelFFC concentrations as indicated (nM) and 12 μM *S. pombe* profilin. C) Representative normalized depolymerization data. Filamentous actin was diluted to 0.1 μM in the presence of increasing concentrations of Delphilin. Capping protein is shown as a control. Analysis of three independent repeats of these assays is shown in Figure 3A. The apparent affinity of mDelFFC for barbed ends is 0.6 nM. D) Representative seeded elongation assays in the presence of increasing concentrations of mDelFC (left) or mDelFCΔ20 (right). The IC_50_ is 7 nM and elongation is ~97% inhibited in both cases, demonstrating that the C-terminus has little influence on the affinity of Delphilin for barbed ends. This value is also similar to that of mDelFFC under the same conditions (IC_50_ = 13 nM). E) Delphilin stimulated actin assembly in the presence of profilin and capping protein. Nucleation is weak but filament elongation demonstrates that Delphilin can protect the barbed end in the presence of capping protein.

## References

Barkó, S., Bugyi, B., Carlier, M.-F., Gombos, R., Matusek, T., Mihály, J., and Nyitrai, M. (2010). Characterization of the Biochemical Properties and Biological Function of the Formin Homology Domains of Drosophila DAAM. J. Biol. Chem. 285, 13154–13169.

Bor, B., Vizcarra, C. L., Phillips, M. L., and Quinlan, M. E. (2012). Autoinhibition of the formin Cappuccino in the absence of canonical autoinhibitory domains. Mol. Biol. Cell 23, 3801–3813.

Courtemanche, N., and Pollard, T. D. (2012). Determinants of Formin Homology 1 (FH1) domain function in actin filament elongation by formins. J. Biol. Chem. 287, 7812–7820.

Dutta, P., Das, S., and Maiti, S. Non Diaphanous Formin Delphilin Acts as a Barbed End Capping Protein. Exp. Cell Res., epub 2017, doi: 10.1016.

Goode, B. L., and Eck, M. J. (2007). Mechanism and function of formins in the control of actin assembly. Annu. Rev. Biochem. 76, 593–627.

Gould, C. J., Maiti, S., Michelot, A., Graziano, B. R., Blanchoin, L., and Goode, B. L. (2011). The Formin DAD Domain Plays Dual Roles in Autoinhibition and Actin Nucleation. Curr. Biol. 21, 384–390.

Harris, E. S., Rouiller, I., Hanein, D., and Higgs, H. N. (2006). Mechanistic differences in actin bundling activity of two mammalian formins, FRL1 and mDia2. J Biol Chem 281, 14383–14392.

Harris, K. M., Jensen, F. E., and Tsao ’, B. (1992). Three-Dimensional Structure of Dendritic Spines and Synapses in Rat Hippocampus (CAI) at Postnatal Day 15 and Adult Ages: Implications for the Maturation of Synaptic Physiology and Long-term Potentiation. J. Neurosci. 12, 2665–2705.

Heimsath, E. G., Jr, and Higgs, H. N. (2012). The C terminus of formin FMNL3 accelerates actin polymerization and contains a WH2 domain-like sequence that binds both monomers and filament barbed ends. J. Biol. Chem. 287, 3087–3098.

Henty-Ridilla, J. L., Rankova, A., Eskin, J. A., Kenny, K., and Goode, B. L. (2016). Accelerated actin filament polymerization from microtubule plus ends. Science 352, 1004–1009.

Higashi, T., Ikeda, T., Murakami, T., Shirakawa, R., Kawato, M., Okawa, K., Furuse, M., Kimura, T., Kita, T., and Horiuchi, H. (2010). Flightless-I (Fli-I) Regulates the Actin Assembly Activity of Diaphanous-related Formins (DRFs) Daam1 and mDia1 in Cooperation with Active Rho GTPase. J. Biol. Chem. 285, 16231–16238.

Higgs, H. N., and Peterson, K. J. (2005). Phylogenetic analysis of the formin homology 2 domain. Mol Biol Cell 16, 1–13.

Hlushchenko, I., Koskinen, M., and Hotulainen, P. (2016). Dendritic spine actin dynamics in neuronal maturation and synaptic plasticity. Cytoskeleton 73, 435–441.

Hotulainen, P., Llano, O., Smirnov, S., Tanhuanpää, K., Faix, J., Rivera, C., and Lappalainen, P. (2009). Defining mechanisms of actin polymerization and depolymerization during dendritic spine morphogenesis. J. Cell Biol. 185, 323–339.

Kashiwabuchi, N. et al. (1995). Impairment of motor coordination, Purkinje cell synapse formation, and cerebellar long-term depression in GluRδ2 mutant mice. Cell 81, 245–252.

Kim, E., Niethammer, M., Rothschild, A., Nung Jan, Y., and Sheng, M. (1995). Clustering of Shaker-type K+ channels by interaction with a family of membrane-associated guanylate kinases. Nature 378, 85–88.

Kohda, K., Kakegawa, W., Matsuda, S., Nakagami, R., Kakiya, N., and Yuzaki, M. (2007). The extreme C-terminus of GluRδ2 is essential for induction of long-term depression in cerebellar slices. Eur. J. Neurosci. 25, 1357–1362.

Kovar, D. R., Harris, E. S., Mahaffy, R., Higgs, H. N., and Pollard, T. D. (2006). Control of the Assembly of ATP- and ADP-Actin by Formins and Profilin. Cell 124, 423–435.

Kovar, D. R., Kuhn, J. R., Tichy, A. L., and Pollard, T. D. (2003). The fission yeast cytokinesis formin Cdc12p is a barbed end actin filament capping protein gated by profilin. J Cell Biol 161, 875–887.

Kupi, T., Gróf, P., Nyitrai, M., and Belágyi, J. (2013). Interaction of formin FH2 with skeletal muscle actin. EPR and DSC studies. Eur. Biophys. J. EBJ 42, 757–765.

Lei, W., Omotade, O. F., Myers, K. R., and Zheng, J. Q. (2016). Actin cytoskeleton in dendritic spine development and plasticity. Curr. Opin. Neurobiol. 39, 86–92.

Li, D. et al. (2011). Dishevelled-associated activator of morphogenesis 1 (Daam1) is required for heart morphogenesis. Dev. Camb. Engl. 138, 303–315.

Li, F., and Higgs, H. N. (2003). The mouse Formin mDia1 is a potent actin nucleation factor regulated by autoinhibition. Curr Biol 13, 1335–1340.

Li, F., and Higgs, H. N. (2005). Dissecting requirements for auto-inhibition of actin nucleation by the formin, mDia1. J Biol Chem 280, 6986–6992.

MacLean-Fletcher, S., and Pollard, T. D. (1980). Identification of a factor in conventional muscle actin preparations which inhibits actin filament self-association. Biochem Biophys Res Commun 96, 18–27.

Matsuda, K., Matsuda, S., Gladding, C. M., and Yuzaki, M. (2006). Characterization of the δ2 Glutamate Receptor-binding Protein Delphilin Splicing Variants With Differential Palmitoylation And An Additional Pdz Domain. J. Biol. Chem. 281, 25577–25587.

McCullough, B. R. et al. (2011). Cofilin-Linked Changes in Actin Filament Flexibility Promote Severing. Biophys. J. 101, 151–159.

Michelot, A., Guerin, C., Huang, S., Ingouff, M., Richard, S., Rodiuc, N., Staiger, C. J., and Blanchoin, L. (2005). The formin homology 1 domain modulates the actin nucleation and bundling activity of Arabidopsis FORMIN1. Plant Cell 17, 2296–2313.

Miyagi, Y. et al. (2002). Delphilin: a Novel PDZ and Formin Homology Domain-Containing Protein that Synaptically Colocalizes and Interacts with Glutamate Receptor δ2 Subunit. J. Neurosci. 22, 803–814.

Moseley, J. B., Sagot, I., Manning, A. L., Xu, Y., Eck, M. J., Pellman, D., and Goode, B. L. (2004). A conserved mechanism for Bni1- and mDia1-induced actin assembly and dual regulation of Bni1 by Bud6 and profilin. Mol Biol Cell 15, 896–907.

Nimchinsky, E. A., Sabatini, B. L., and Svoboda, K. (2002). Structure and Function of Dendritic Spines. Annu. Rev. Physiol. 64, 313–353.

Otomo, T., Otomo, C., Tomchick, D. R., Machius, M., and Rosen, M. K. (2005). Structural basis of Rho GTPase-mediated activation of the formin mDia1. Mol Cell 18, 273–281.

Paul, A. S., and Pollard, T. D. (2008). The role of the FH1 domain and profilin in formin-mediated actin-filament elongation and nucleation. Curr Biol 18, 9–19.

Penzes, P., Cahill, M. E., Jones, K. A., VanLeeuwen, J.-E., and Woolfrey, K. M. (2011). Dendritic spine pathology in neuropsychiatric disorders. Nat. Neurosci. 14, 285–293.

Peters, A., and Kaiserman-Abramof, I. R. (1970). The small pyramidal neuron of the rat cerebral cortex. The perikaryon, dendrites and spines. Am. J. Anat. 127, 321–355.

Pruyne, D. (2016). Revisiting the Phylogeny of the Animal Formins: Two New Subtypes, Relationships with Multiple Wing Hairs Proteins, and a Lost Human Formin. PLOS ONE 11, e0164067.

Pruyne, D., Evangelista, M., Yang, C., Bi, E., Zigmond, S., Bretscher, A., and Boone, C. (2002). Role of formins in actin assembly: nucleation and barbed-end association. Science 297, 612–615.

Quinlan, M. E., Hilgert, S., Bedrossian, A., Mullins, R. D., and Kerkhoff, E. (2007). Regulatory interactions between two actin nucleators, Spire and Cappuccino. J Cell Biol 179, 117–128.

Ramabhadran, V., Gurel, P. S., and Higgs, H. N. (2012). Mutations to the Formin Homology 2 Domain of INF2 Protein Have Unexpected Effects on Actin Polymerization and Severing. J. Biol. Chem. 287, 34234–34245.

Ramalingam, N., Zhao, H., Breitsprecher, D., Lappalainen, P., Faix, J., and Schleicher, M. (2010). Phospholipids regulate localization and activity of mDia1 formin. Eur J Cell Biol 89, 723–732.

Rosales-Nieves, A. E., Johndrow, J. E., Keller, L. C., Magie, C. R., Pinto-Santini, D. M., and Parkhurst, S. M. (2006). Coordination of microtubule and microfilament dynamics by Drosophila Rho1, Spire and Cappuccino. Nat Cell Biol 8, 367–376.

Rose, R., Weyand, M., Lammers, M., Ishizaki, T., Ahmadian, M. R., and Wittinghofer, A. (2005). Structural and mechanistic insights into the interaction between Rho and mammalian Dia. Nature 435, 513–518.

Rubenstein, P. A. (1990). The functional importance of multiple actin isoforms. BioEssays 12, 309–315.

Salomon, S. N., Haber, M., Murai, K. K., and Dunn, R. J. (2008). Localization of the Diaphanous-related formin Daam1 to neuronal dendrites. Neurosci. Lett. 447, 62–67.

Sato, A., Khadka, D. K., Liu, W., Bharti, R., Runnels, L. W., Dawid, I. B., and Habas, R. (2006). Profilin is an effector for Daam1 in non-canonical Wnt signaling and is required for vertebrate gastrulation. Development 133, 4219–4231.

Schindelin, J. et al. (2012). Fiji: an open-source platform for biological-image analysis. Nat. Methods 9, 676–682.

Schönichen, A. et al. (2013). FHOD1 is a combined actin filament capping and bundling factor that selectively associates with actin arcs and stress fibers. J. Cell Sci. 126, 1891–1901.

Seth, A., Otomo, C., and Rosen, M. K. (2006). Autoinhibition regulates cellular localization and actin assembly activity of the diaphanous-related formins FRLalpha and mDia1. J Cell Biol 174, 701–713.

Smith, M. B., Li, H., Shen, T., Huang, X., Yusuf, E., and Vavylonis, D. (2010). Segmentation and tracking of cytoskeletal filaments using open active contours. Cytoskelet. Hoboken NJ 67, 693–705.

Sreerama, N., and Woody, R. W. (1993). A Self-Consistent Method for the Analysis of Protein Secondary Structure from Circular Dichroism. Anal. Biochem. 209, 32–44.

Takeuchi, T. et al. (2008). Enhancement of Both Long-Term Depression Induction and Optokinetic Response Adaptation in Mice Lacking Delphilin. PLOS ONE 3, e2297.

Vizcarra, C. L., Bor, B., and Quinlan, M. E. (2014). The role of formin tails in actin nucleation, processive elongation, and filament bundling. J. Biol. Chem. 289, 30602–30613.

Vizcarra, C. L., Kreutz, B., Rodal, A. A., Toms, A. V., Lu, J., Zheng, W., Quinlan, M. E., and Eck, M. J. (2011). Structure and function of the interacting domains of Spire and Fmn-family formins. Proc Natl Acad Sci U A 108, 11884–11889.

Walikonis, R. S., Jensen, O. N., Mann, M., Provance, D. W., Mercer, J. A., and Kennedy, M. B. (2000). Identification of proteins in the postsynaptic density fraction by mass spectrometry. J. Neurosci. Off. J. Soc. Neurosci. 20, 4069–4080.

Xu, Y., Moseley, J. B., Sagot, I., Poy, F., Pellman, D., Goode, B. L., and Eck, M. J. (2004). Crystal structures of a Formin Homology-2 domain reveal a tethered dimer architecture. Cell 116, 711–723.

Yamashita, T. et al. (2005). Identification and characterization of a novel Delphilin variant with an alternative N-terminus. Mol. Brain Res. 141, 83–94.

Yoo, H., Roth-Johnson, E. A., Bor, B., and Quinlan, M. E. (2015). Drosophila Cappuccino alleles provide insight into formin mechanism and role in oogenesis. Mol. Biol. Cell 26, 1875–1886.

Zalevsky, J., Grigorova, I., and Mullins, R. D. (2001). Activation of the Arp2/3 complex by the Listeria acta protein. Acta binds two actin monomers and three subunits of the Arp2/3 complex. J Biol Chem 276, 3468–3475.

